# Modelling isoscapes using mixed models

**DOI:** 10.1101/207662

**Authors:** Alexandre Courtiol, François Rousset

## Abstract

**Abstract:** Isoscapes are maps depicting the continuous spatial (and sometimes temporal) variation in isotope composition. They have various applications ranging from the study of isotope circulation in the main earth systems to the determination of the provenance of migratory animals. Isoscapes can be produced from the fit of statistical models to observations originating from a set of discrete locations. Mixed models are powerful tools for drawing inferences from correlated data. While they are widely used to study non-spatial variation, they are often overlooked in spatial analyses. In particular, they have not been used to study the spatial variation of isotope composition. Here, we introduce this statistical framework and illustrate the methodology by building isoscapes of the isotope composition of hydrogen (measured in *δ*^2^H) for precipitation water in Europe. For this example, the approach based on mixed models presents a higher predictive power than a widespread alternative approach. We discuss other advantages offered by mixed models including: the ability to model the residual variance in isotope composition, the quantification of prediction uncertainty, and the simplicity of model comparison and selection using an adequate information criterion: the conditional AIC (cAIC). We provide all source code required for the replication of the results of this paper as a small `R` package to foster a transparent comparison between alternative frameworks used to model isoscapes.

**Abbreviations used in this paper:**

- AIC: Akaike Information Criterion
- BLUP: Best Linear Unbiased Predictor
- BWR: a method for building isoscape introduced by Bowen and Wilkinson (2002) and Bowen and Revenaugh (2003)
- cAIC: conditional Akaike Information Criterion
- DHGLM: Double Hierarchical Generalised Linear Model
- GLM: Generalised Linear Model
- GLMM: Generalised Linear Mixed-effects Model
- GNIP: Global Network for Isotopes in Precipitation
- LM: Linear Model
- LMM: Linear Mixed-effects Model
- MAE: Mean Absolute Error
- ML: Maximum Likelihood
- REML: Restricted Maximum Likelihood
- RMSE: Root Mean Squared Error

## 1 INTRODUCTION

In this paper, we introduce a methodology based on mixed models for describing how the ratio of light to heavy isotopes found in environmental materials varies in space. The spatial variation in isotope composition is typically represented by maps called isoscapes (from **iso**topic land**scapes**). Isoscapes have multiple purposes which have be reviewed by Bowen (2010) and West et al. (2010, chapters 13-20). Briefly, they are used to study the relationship between environmental (biological, climatological, geological and hydrological) factors and the isotope composition, which provides information on the major earth systems such as the cycle of carbon, nitrogen or water. Isoscapes are also increasingly used in ecology and forensic science to determine where organisms (or part of) are coming from; a task that is possible because the isotope composition of living organisms is influenced by that of their environment.

Similarly to what is usually done to compute weather forecasts, the physicochemical processes known to impact isotope composition can be simulated numerically to obtain isoscapes (Noone and Sturm, 2010). These dynamical models are useful because they allow for the precise study of how the different processes interplay to shape the spatial distribution of isotopes. However, the predictive power of such models is currently too low for many applications. The alternative approach consists in ignoring the precise mechanistic details shaping the spatial distribution of isotopes and seeing isoscape construction as a statistical prediction problem: from a limited number of observations at fixed spatial locations, the goal is to predict the continuous spatial process that generates the data.

Within the scope of the statistical approach, a widely used method to model isoscapes is that of Bowen and colleagues (Bowen and Revenaugh, 2003; Bowen and Wilkinson, 2002), but other approaches have been considered. Often the models are fitted in two steps: a regression model to environmental variables is fitted first, and then a spatial model is fitted to the residuals of this first fit. This second fit may use traditional *Kriging* techniques that are broadly known in spatial statistics (Cressie, 1993; Zimmerman and Stein, 2010), but various other techniques have been considered in the isoscape literature (*e.g.* Bowen and Revenaugh, 2003). Several authors have argued that Kriging is best understood as prediction under a particular linear mixed model (not to be confused with mixing models) (*e.g.* Christensen, 1991, 2011; Diggle and Ribeiro, 2007; Diggle et al., 1998; Pollice and Bilancia, 2007), and therefore mixed model methods could be used to model isoscapes.

Fitting isoscapes using mixed models should present several benefits. First, mixed models fulfil the needs covered by traditional Kriging methods, but contrary to traditional Kriging approaches mixed models are based on clear and explicit statistical definitions (Zimmerman and Stein, 2010). Second, mixed models offer the possibility of covering a wider range of assumptions than usually considered in the Kriging and isoscape literature. Third, mixed-model theory provides likelihood-based methods to fit jointly the spatial and environmental components of a model, which is generally known to reduce biases in parameter estimates (Cressie, 1993). Fourth, the likelihood framework makes this approach superior in terms of inference possibilities as well as for model comparison or selection procedures. For example, after a single fitting procedure mixed models provide confidence intervals, prediction intervals, and model selection criteria. Last but not least, mixed models are widely used to study many kinds of spatial but also non-spatial problems (Bolker et al., 2009; Pinheiro et al., 2007; Venables and Ripley, 2002); therefore, more flexible models keep being proposed (*e.g.* Lee and Nelder, 2006), efficient algorithms to fit these models are implemented in both general and dedicated statistical software, and guidance on how to use mixed models keeps accumulating (Bolker et al., 2009).

Despite the benefits of the mixed-models based approach, it is not routinely used to analyse the spatial distribution of isotopes. Early works relied on so-called ordinary Kriging (*e.g.* Gutierrez, 1991), which does not estimate the effects of environmental variables. Later on, Bowen and Wilkinson (2002) and Bowen and Revenaugh (2003) introduced such estimation. Albeit being model-based and representing an improvement over previous methods, the statistical approaches used by Bowen and colleagues do not use Kriging methods (and mixed models *a fortiori*). Kriging has been reintroduced in later works (*e.g.* Terzer et al., 2013; Van der Veer et al., 2009), but the Kriging is done without using mixed models explicitly.

Here, we explain how to build isoscapes using mixed models. Our methodology is general and can handle different datasets, isotopes, or environmental materials. As a study case, we will build the isoscape for the hydrogen composition (measured in *δ*^2^H) of precipitation water using data from the Global Network for Isotopes in Precipitation (IAEA/WMO, 2017). This will illustrate how to fit the mixed models underlying isoscapes, how to build different isoscapes from the fitted models, how to weight monthly predictions by precipitation amounts to obtain the traditional annual weighted predictions, and how to compare different statistical models fitted on the same data. We will also show that, at least for the data we use, our approach is more accurate than the widely used approach from Bowen and Wilkinson (2002) and Bowen and Revenaugh (2003). Finally, we will discuss the benefits and limits of the mixed-model based approach with respect to practical applications.

We implement the analysis in the free open-source statistical software `R` (R Core Team, 2017) and perform most computations using the package `spaMM` (Rousset and Ferdy, 2014). This package is a multi-purpose package that allows the user to fit mixed models with (but also without) spatial autocorrelation by maximum (or restricted maximum) likelihood. We provide all the source code necessary to produce all the figures and tables present in this paper under the form of a small package (available in Supplementary material). We developed this package for reproducibility purposes only, and we did not design this package for general usage on other datasets or applications. For general usage we recommend instead to either use `spaMM` directly, or to use the `IsoriX` package which is a wrapper for `spaMM` specifically developed for modeling the spatial distribution of isotopes, as well as to assign organisms to their origin location (Courtiol et al., 2016). We assume little prior knowledge of mixed models in this paper and we define some key concepts related to these models in the glossary. We assume nonetheless knowledge of the basic concepts generally introduced in the context of generalized linear models (GLMs) such as linear predictor, regression coefficients, or residuals; otherwise see Venables and Ripley (2002, Chapter 7) for an introduction to GLM and Diggle et al. (2003) for a gentle introduction to spatial mixed models.

## 2 METHODOLOGY

### 2.1 Defining the spatial variation in isotope composition with mixed models

Statistically, the spatial variation in the isotope composition of an environmental sample can be expressed as a linear mixed-effects model (LMM, see glossary) that describes, for each observation *i* at a geographical location *g* (a given monitoring station where the precipitation water is collected), the value of the response variable `iso`_*gi*_ (a given measurement of the isotope composition) as

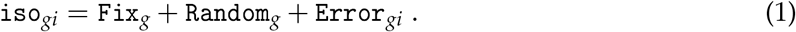

Hereafter, we denote this model as the *mean model*. The terms `Fix`, `Random` and `Error` denote the fixed effects, the random effects, and the residual errors, respectively (see glossary). By definition, `Error`_*gi*_ is independent for each observation *i*, which distinguishes it from the effect `Random`_*g*_ which is common to all observations at the location *g*. We will now detail each of these three terms.

Let us first consider the fixed effects (`Fix`). Different parameterizations have been proposed to model how spatial and climatic covariates influence the isotope composition of hydrogen and oxygen in precipitation water (*e.g.* Bowen and Wilkinson, 2002; Terzer et al., 2013; Van der Veer et al., 2009). For now, we consider the following parametrization (but we will consider others below):

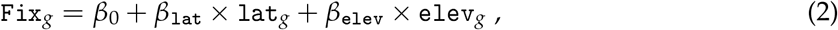

where the two covariates involved are `lat`_*g*_, which represents the latitude of the location *g*, and `elev`_*g*_, which represents its elevation. We use the latitude^1^ and the elevation to approximate the effects that temperature exerts on the isotope composition in precipitation water (Bowen and Wilkinson, 2002). Contrary to temperature *per se*, the indirect proxies we choose (latitude and elevation) are readily available for any location, which will be crucial to predict the isotope composition over a geographic area. The regression coefficient *β*_0_ represents an intercept, that is the predicted mean isotopic values at sea level (`elev` = 0) at the equator (`lat` = 0). The regression coefficients *β*_lat_ and *β*_elev_ transform the latitude and the elevation into isotope compositions.

Second, we consider random effects in the model (`Random`) in order to capture two sources of variation accounting for departure from the mean composition in isotopic values predicted by the fixed effects:

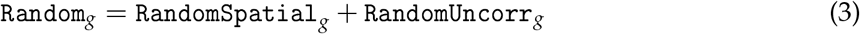

The source of variation `RandomSpatial` stems from the many unmeasured environmental factors that vary spatially but that are not (and could not all be) considered as fixed effects. Such factors are sources of spatial autocorrelation: nearby locations are often traversed by the same air masses and are thus more similar in their isotope composition than distant locations. We can describe this correlation between the random effects in different locations as a function of their geographical distance.

To represent this spatial autocorrelation, we use the so-called Matérn correlation function (Matérn, 1960). Specifically, we assume that the correlation between random effects at spatial distance *d*_*ab*_ between two given location (*a* and *b*) is of the form *M*_*v*_ (*ρd*_*ab*_) where *ρ* is a spatial scale parameter and *M*_*v*_ is the Matérn correlation function, which can be written as:

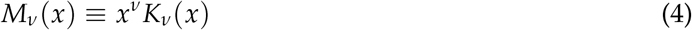

where *K*_*v*_ is the Bessel function of second kind and order *v*, and *v* is the “smoothness” parameter (the higher *v* is, the smoother are the realized surfaces at a small scale). The Matérn correlation function is appropriate to fit autocorrelated processes with more or less rugged realizations, and is the most useful correlation model for a wide range of applications (Stein, 1999; see also *e.g.* Diggle and Ribeiro, 2007). It includes the commonly used exponential and squared exponential (or “Gaussian“) correlation functions as special cases (for *v* = 0.5 and *v* → ∞ to, respectively). Any *v* > 0 may be considered in Euclidean spaces of any dimension. However, Euclidean distances are poor approximations of geographical distances over large spatial ranges. We thus recommend the use of an appropriate distance method such as orthodromic distances to account for the earth curvature. On a sphere, 0 < *v* ≤ 0.5, as for higher values, matrices of correlation values between different locations may not be positive semi-definite and therefore invalid as correlation matrices.

Thus, *v* = 0.5 is the highest value that can be considered for isoscapes with orthodromic distances.

We therefore assume that the random effects in each location (*i.e.* `RandomSpatial`_*g*_) are normally distributed, with mean zero, variance λ_RS_, and correlation

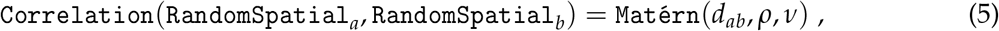

where *ρ* is a scaling factor for the distance, and *v* is the smoothness parameter of the Matérn.

The second source of variation - `RandomUncorr` - represents factors that differ between locations but that are not spatially correlated. Examples could be weather conditions (micro-climates) or sources of measurement error that vary locally. This second random effect has mean zero, variance denoted λ_RU_, takes the same value for all realizations drawn from the same location, and is uncorrelated between different locations.

Finally, the residual term of the model (`Error`) is, by definition for LMMs, normally distributed with mean zero and a variance *ϕ*_*g*_ This variance *ϕ*_*g*_ is typically assumed constant in LMMs. However, as we shall see, the per-location estimates of the variation between observations `variso`_*g*_ are themselves strongly spatially structured. We thus propose to model the square of `Error`_*gi*_ as a random variable with expectation:

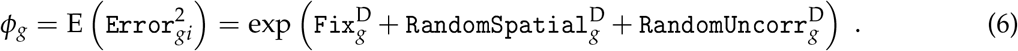

Hereafter, we denote this model as the *residual dispersion model*. In this model, we consider that the fixed-effects component `Fix`^D^ is reduced to an intercept 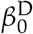. The spatial random effect (`RandomSpatial`^D^) is again normally distributed with zero mean, but has now a variance 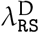 and a Matérn correlation structure with parameters *ρ*^D^ and *v*^D^. The uncorrelated random effect (`RandomUncorr`^D^) is also normally distributed with zero mean, has now a variance 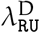 and is again uncorrelated between different locations. Note that the roman D index is not a mathematical exponent: it denotes parameters of the residual dispersion model. Note that there is no particular reason to expect the estimatations for the two random effects of the residual dispersion model to equate those for the two random effects of the mean model. We will thus have to estimate four different random effects. The exponential in eq. 6 ensures that the predicted value of the variance *ϕ*_*g*_ in any location will be positive.

### 2.2 The dataset

For demonstration, we will fit the aforementioned mixed models on monthly measurements of the isotope composition of the hydrogen (technically expressed as *δ*^2^H) found in precipitation water. The dataset was provided by the Global Network for Isotopes in Precipitation (IAEA/WMO, 2017), which distributes freely isotope measurements from precipitation water collected all over the world. The coverage of the GNIP data is variable depending on the world region, and we will restrict our analysis to Europe (here defined as locations within the latitude 30°00′N - 70°00′N and the longitude 30°00′W - 60°00′E), which is a particularly data dense area. We discarded observations for which required information was missing, those corresponding to a month and monitoring station only observed a single time, and those for which “month” samples have been derived from a precipitation collection based on 25 days or less, or on 35 days or more. We also excluded the data from one monitoring station for which sampled names indicated they were half month samples instead of monthly ones. After this filtering procedure, our data yielded a total of 30546 monthly measurements associated with 326 different locations. These data cover 658 out of the 660 possible months covering the study period (1960-2014). The raw data provided by the GNIP project are readily aggregated into monthly measurements for each available year and sampling location. We chose to aggregate these data further, across years, so to obtain a single mean and variance estimate for each available month-location combination. We aggregated the data this way for faster fitting of both the mean model and the residual dispersion model, as detailed below. We denote the final aggregated dataset `GNIPdataEU` (table 1).

**Table 1.** First 15 rows of the `GNIPdataEU` dataset. We derived this dataset from data provided by the Global Network of Isotopes in Precipitation (IAEA/WMO, 2017). It contains a total of 3313 rows and 8 columns. These columns provide the month of the sample collection (`month`), the mean and variance of *δ*^2^H measured across years for the month and locality considered (`meaniso`, `variso`), the number of years considered (`nyear`), the World Meteorological Organization code of the monitoring station where the water sample has been collected (`stationID`), and the approximate coordinates and elevation of the sampling site as provided by GNIP (`lat`, `long`, `elev`).

| month | meaniso | variso | nyear | stationID | lat | long | elev |
| --- | --- | --- | --- | --- | --- | --- | --- |
| 1 | -49.68 | 69.16 | 10 | 142700 | 58.10 | 6.57 | 13 |
| 2 | -48.86 | 202.83 | 10 | 142700 | 58.10 | 6.57 | 13 |
| 3 | -46.36 | 89.24 | 9 | 142700 | 58.10 | 6.57 | 13 |
| 4 | -45.83 | 104.17 | 7 | 142700 | 58.10 | 6.57 | 13 |
| 5 | -43.60 | 113.02 | 10 | 142700 | 58.10 | 6.57 | 13 |
| 6 | -34.16 | 60.38 | 10 | 142700 | 58.10 | 6.57 | 13 |
| 7 | -35.26 | 220.08 | 9 | 142700 | 58.10 | 6.57 | 13 |
| 8 | -42.80 | 152.14 | 8 | 142700 | 58.10 | 6.57 | 13 |
| 9 | -52.76 | 211.23 | 8 | 142700 | 58.10 | 6.57 | 13 |
| 10 | -49.01 | 195.41 | 9 | 142700 | 58.10 | 6.57 | 13 |
| 11 | -52.01 | 284.18 | 10 | 142700 | 58.10 | 6.57 | 13 |
| 12 | -54.53 | 379.32 | 9 | 142700 | 58.10 | 6.57 | 13 |
| 1 | -160.75 | 311.84 | 6 | 206000 | 68.68 | 21.53 | 403 |
| 2 | -142.87 | 317.88 | 6 | 206000 | 68.68 | 21.53 | 403 |
| 3 | -146.82 | 76.54 | 6 | 206000 | 68.68 | 21.53 | 403 |

### 2.3 Fitting approach

We now want to fit the mixed models on the `GNIPdataEU` dataset to obtain estimates for the fixed effects *β*_0_, *β*_lat_, *β*_elev_ and the random effect parameters λ_RS_, *ρ*, *v*, λ_RU_ of the mean model (eq. 1), and likewise the estimates for the fixed and random effect parameters 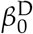, and 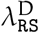, *ρ*^D^, *v*^D^, 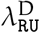 of the residual dispersion model (eq. 6). Because the spatial variation in the hydrogen isotope composition in precipitation water varies seasonally, we will fit both models on each of the twelve months independently. We will thus use twelve different subsets of `GNIPdataEU` corresponding to each month (`GNIPdataEU1` to `GNIPdataEU12`). The following discussion explains how to fit both models for a given month and we choose January for this illustration (*i.e.* we use `GNIPdataEUl`). We will later show how model fits can be combined across months to produce annual averaged predictions.

The procedure consists in fitting first the residual dispersion model to the observed variances in each location, before fitting the mean model. In this procedure, the fixed effects and random effects parameters are estimated jointly for the mean model (but not for both models simultaneously). Our estimating procedure is thus an iterative full-likelihood fit of the mean observations, and it allows for full likelihood-based inference under the mean model, given the estimates of the parameters of the residual dispersion model.

#### 2.3.1 Fitting the residual dispersion model

We first use the spatial mixed-effect model for the expectation of the residuals of the mean model (eq. 6), and fit this residual dispersion model to the variance estimates `variso`, derived from `nyear`_g_ monthly observations in each location *g* (see table 1). From our earlier assumption that `Error`_gi_ is normally distributed with variance *ϕ*_*g*_, `variso` is Gamma-distributed with mean *ϕ*_*g*_ and variance 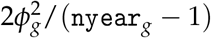.^2^ We can therefore model variso using a Gamma generalized linear mixed-effects model (Gamma GLMM).

To account for the variance 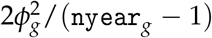 of `variso` in the residual dispersion model, we need to consider how residual variance is represented in a Gamma GLMM. The package `spaMM` follows the established usage for Gamma GLMs (McCullagh and Nelder, 1989), where the Gamma-distributed response is by default parametrized by its mean *μ* and a dispersion parameter Φ such that the variance of the response is *μ*^2^Φ. In addition, known prior weights *w*_*i*_ can be specified, such that the dispersion for each level *i* of the response variable is Φ_*i*_ = Φ/*w*_*i*_. We can therefore represent 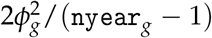 as *μ*^2^*ϕ*^D^/*w*_*g*_ for *μ* = *ϕ*_*g*_, *ϕ*^D^ = 2, and prior weights *w*_*g*_ = `nyear`_*g*_ − 1 for each location *g*.

We now use this result to fit the residual dispersion model on the data for January using the function from `fitme()`, from `spaMM` as follow:

~~~
dispfit1 <- fitme(
  formula = variso ^~^ 1 + Matern(1|long + lat) + (1|stationID),
  family = Gamma(link = log), data = GNIPdataEU1, fixed = list(phi = 2),
  prior.weights = nyear - 1, control.dist = list(dist.method = "Earth"),
  method = "REML")
~~~

This simple call produces `R` an object denoted that `dispfit1` contains estimates of the parameters of the residual dispersion model. The arguments `formula` and `family` are used to specify the residual dispersion model. Specifically, in the `formula`, the “1” indicates to fit one intercept, the term `Matérn (1|long + lat)` indicates to fit a spatially-structured random effect following a Matérn correlation structure where the variable `long` and `lat`^3^ will be used for the computation of the distance between observations, and the term `(1|stationID)` indicates to fit an uncorrelated random effect whose realizations will be identical for all observations with the same value for the variable `stationID`. The argument `family` specifies the Gamma family with log link, where the link corresponds to the exponential in eq. 6. The argument `data` indicates in which `R` object the dataset is stored. The argument `fixed` is used to fix parameter values(s) in the model, and we use it here specifically to fix the residual dispersion *ϕ*^D^ = 2 in `dispfitl`. The argument `prior.weight` is used to provide the prior weights. The argument `control.dist` specifies the `Earth` computation method for orthodromic distances, instead of the default distance method in `spaMM`(Euclidean distances). Finally, the argument `method` specifies that restricted maximum likelihood is used to fit the model. For non-Gaussian GLMMs, the REML `method` actually implements an approximate concept of restricted likelihood defined by Lee et al. (2006, p. 187). We recommend to stick to REML because other forms of maximum likelihood lead to biases in estimation of prediction variances, which we will use to build the isoscapes.

#### 2.3.2 Fitting the mean model

The second step of our procedure is to fit the mean model. This is a LMM with two distinctive features. First, we need the predictions 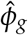 of the expected square of the residual error in each location, given by the fit of the residual dispersion model. We can derive the predictions from `dispfit1` with the function `predict()` from `spaMM` and store them within our working dataset with the call:

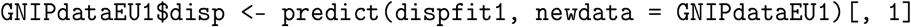

Second, since the fit of the main model is a fit to the mean of `nyear`_*g*_ observations in each location *g*, we will also specify that the residual variance for each such mean is 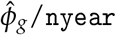 by using prior weights `nyear`.

We fit the mean model for January using the function `fitme()` as before:

~~~
meanfit1 <- fitme(
  formula = meaniso ^~^ 1 + lat + elev + Matern(1|long + lat) + (1|stationID),
  family = gaussian(link = identity), data = GNIPdataEU1,
  resid.model = list(formula= ^~^ 0 + offset(disp), family = Gamma(link = identity)),
  prior.weights = nyear, control.dist = list(dist.method = "Earth"),
  method = "REML")
~~~

This call produces an object denoted `meanfit1` that contains estimates of the parameters of the
mean model. Here, we specified explicitly the form that residual errors take using the argument `resid.model`. Its `formula` element forces the residual variance to be exactly as predicted by `dispfit1`, by way of the standard `offset()` syntax, combined with the 0 which indicates no to fit an intercept. The model for residual errors expected by `fitme` must belong to the Gamma family, which we specify using the `family` argument. The default link for the Gamma family being `log`, the exponential of the offset would serve as prediction for *ϕ*_*g*_ In order to consider the offset directly, we must thus also specify that the fitting procedure must use the non-default identity link.

### 2.4 Building isoscapes

We will use the fits obtained above to predict the spatial variation in hydrogen isotope composition for an average month of January during 1960-2014. Specifically, we aim to predict three different isoscapes: the traditional isoscape that represents the prediction for *δ*^2^H in each location (called point prediction), and two additional maps representing the prediction variance and the residual variance.

The prediction variance has the following meaning. Our model assumes that that there is a true unknown isoscape, which is fixed (in particular, it does not change as data accumulate) but which is represented by the mixed model as a random draw from possible realizations of isoscapes (random draws of the Matérn-correlated process and of any other random effect). We infer this realized isoscape by fitting the model to a limited amount of data, with some uncertainty since different random draws of the unknown isoscape may give the same observed data. There is thus a conditional distribution of possible true isoscapes given the data. For LMM, the Best Linear Unbiased Prediction (BLUP) is the mean of this conditional distribution (Robinson, 1991), and the prediction variance is ideally the mean squared difference between the true unknown value of the linear predictor and the BLUP at a given location. Put more simply, the prediction variance quantifies how precise our point predictions are.

The residual variance has a different meaning. It estimates the variance of new observations drawn from the true unknown isoscape at a given location. This variance corresponds to the point prediction from the residual dispersion model. It thus represents here the between-year variance in each location.

#### 2.4.1 The dataset for predictions

Because it is not possible to make prediction for an infinite number of locations, we will predict the isotope composition on a regular array within the extent of latitudes and longitudes of our selected area. We call these locations *prediction locations* The prediction locations are thus arranged in rows of equal latitudes and columns of equal longitudes forming a regular geographic array or *rasters*.

To compute a prediction for each prediction location, we need the parameter estimates from the fit of the mean model and from the fit of the residual dispersion model provided by `meanfit1` and `dispfit1`, respectively. The objects `meanfit1` and `dispfit1` also contain all necessary parameters required to compute the prediction and residual variances. For each prediction location, we also need the values for all covariates used in the mixed models. In the case of the models presented above, the covariates needed are the latitude, longitude and elevation. The latitude and longitude is directly defined by the prediction location. Other covariates (the elevation in our case) also come from rasters. Therefore, the maximal resolution one can achieve to predict isoscape values are constrained by the lower resolution of the required rasters. Here, we use a raster called `elevraster`, which stores elevations for the entire Earth at a resolution of approximately one elevation value per 100 square-km. This resolution thus defines the resolution at which we compute the predictions forming the isoscapes.

To prepare all the values for the covariates needed for the prediction procedure, we first load the elevation raster and crop it to the extent of Europe. Then, we extract the geographic coordinates of each elevation from this object to build a table (`a data.frame,` in `R` language) that contains all the values for the covariates needed for the prediction procedure. This can all be done easily `R` in with the help of functions from the packages `raster` and `sp` (see Supplementary material for `R` code). We call `datapred` the table containing the values for the covariates needed for the prediction procedure (table 2).

**Table 2.** First six rows of the object `datapred`. The columns `long`, `lat`, `elev` and `stationID` provide the covariate values for predictions. Notice that in the column `stationID`, we gave a new unique name for each prediction location. This enables `spaMM` to assign to the new locations the expected realization of the uncorrelated random effect (*i.e.* zero) instead of the value of the uncorrelated random effect inferred for locations where monitoring stations exist.

| lat | long | elev | stationID |
| --- | --- | --- | --- |
| 69.42 | -29.58 | 2136.03 | new1 |
| 69.42 | -28.75 | 2163.40 | new2 |
| 69.42 | -27.92 | 2081.96 | new3 |
| 69.42 | -27.08 | 1787.53 | new4 |
| 69.42 | -26.25 | 1437.74 | new5 |
| 69.42 | -25.42 | 798.55 | new6 |

#### 2.4.2 Predicting the isoscapes for January

Using the table `datapred` and the parameter estimates obtained from `dispfit1`, we start by predicting the residual dispersion variance for each prediction location, according to

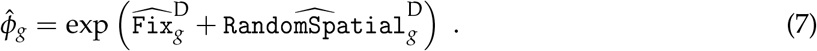

Hat symbols denote estimations of fixed effects, predictions of random effects, and functions of them. On the right-hand side of the equation, they thus denote estimations and predictions from the fit of the dispersion model. Note that the uncorrelated random effect is not apparent because the prediction of an uncorrelated random effect is necessarily zero for any new location.

This computation is done by a first call to the `spaMM` function `predict()`. We directly store the predictions for the residual variance in `datapred`. In `R` language, this gives:

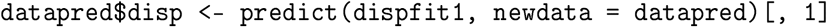

We can then use the table `datapred` and the parameter estimates obtained from `meanfit1` to compute the predicted isotope value over all possible observations *i* at a location *g* as

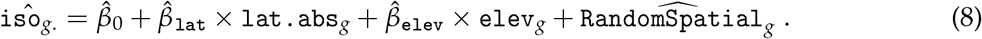

Again, the uncorrelated random effect is not apparent because its prediction is zero for any new locations 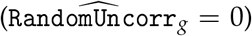. This computation is done by a second call to the `spaMm` function `predict():`

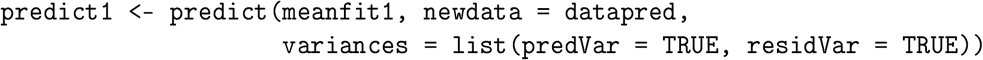

In addition to the point prediction for each prediction location, this second call to `predict` uses the information obtained from the first call (*i.e.* `datapred$disp`) to compute the prediction variance of the mean model.

We finally extract the point predictions, the prediction variance and the residual variance, and store them within a single `R` object denoted `pred1` which also contains the prediction locations (see table 3 for content):

~~~
pred1 <- datapred[, c("lat", "long")]
pred1$pred <- predict1[, 1]
pred1$predVar <- attr(predict1, "predVar")
pred1$residVar <- attr(predict1, "residVar")
~~~

**Table 3.** First six rows of the object `predl`, which contains the predictions for the month of January. The columns provide the coordinates of the prediction locations, *i.e.* latitude (`lat`) & longitude (`long`), as well as the predicted isotope composition (`pred`), the prediction variance (`predvar`) and the residual variance (`residvar`) associated to these locations.

| lat | long | pred | predVar | residVar |
| --- | --- | --- | --- | --- |
| 69.42 | -29.58 | -123.10 | 396.24 | 166.96 |
| 69.42 | -28.75 | -123.41 | 387.81 | 167.39 |
| 69.42 | -27.92 | -122.32 | 378.89 | 167.83 |
| 69.42 | -27.08 | -118.45 | 368.83 | 168.29 |
| 69.42 | -26.25 | -113.88 | 359.37 | 168.76 |
| 69.42 | -25.42 | -105.52 | 349.92 | 169.24 |

Once we have the point prediction, the prediction variance and residual variance for each prediction location, we can then turn the isoscape data we want to represent (`pred`, `predVar`, or `residvar`) into a raster and plot the raster for visualization using `R` an function dedicated for plotting rasters, such as `levelplot()` from the package `rasterVis` (see Supplementary material for `R` code). We will show these plots as well as plots corresponding to the following section in *Results*.

#### 2.4.3 Predicting the annual isoscapes

We obtain a total of 12 pairs of fitted mixed models by repeating the fitting procedure of the mean model and of the residual dispersion model on the subsets of `GNIPdataEU` corresponding to all months of a year (`GNIPdataEUl` to `GNIPdataEU12`). Each pair of mixed models can readily be used to produce the point prediction, prediction variance, or residual variance, for a given month as we have just explained it for January. This leads to the construction of tables to `pred1`, to `pred12`, analogously to the one shown in table 3. Then the isoscapes for each month can easily be derived from these tables, as mentioned above.

We can also use all 12 obtained tables to compute the annual averaged point prediction, prediction variance and residual variance for each prediction location. Under the assumption that the observations collected for each month are independent, at each location the point prediction for a complete year is simply the average of the 12 point predictions (one for each month). Accordingly, at each prediction location the prediction variance for the annual average becomes the sum of the prediction variances for the 12 months divided by 12^2^ and similarly the residual variance for the annual average becomes the sum of the 12 residual variances divided by 12^2^.

It is also possible to assign a different weight to each monthly isoscape before combining them. As an illustration, we will compute the annual average weighted by the estimated amount of precipitation that fell in each month at each prediction location. We chose this example because the use of such annual average weighted by precipitation amounts is common in the isoscape literature (Bowen, 2010). For this we need to derive weight for each month from precipitation data at each prediction location. These precipitation data can be obtained freely from WorldClim 1.4 (Hijmans et al., 2005), which provides such information as a set of 12 rasters (one for each month) at a resolution of ca. one value per square kilometre. After downloading these rasters, we use the function `extract()` from the package `raster` to extract the precipitation amount for the 12 months at each prediction location. Then, if we call `precip`_*m,g*_ the precipitation amount for a month *m* at location *g*, the weight *w*_*m,g*_ for the month m is directly the relative precipitation during this month, that is 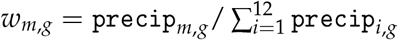. We finally obtain the point prediction of the precipitation-amount weighted annual average as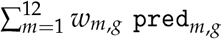, the prediction variance as 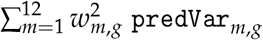 and the corresponding residual variance as 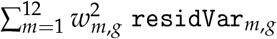

Other work have produced isoscapes of (precipitation-amount weighted) annual averages differently. Instead of combining predictions per month as we propose, the common approach is to first compute annual averages directly on the data and then fit the model on the averaged data. We chose a different approach because we wanted to avoid two main limitations characterizing such an alternative approach. First, the traditional approach implies to only retain years for which all (or most) months have been sampled. This would imply to discard many data from our dataset and thus information would be lost. Second, the traditional approach does not illustrate temporal variation underlying the annual pattern, while our approach allows the creation of different isoscapes for each month.

### 2.5 Model comparison and model selection

In contrast to mechanistic dynamical models (Noone and Sturm, 2010), the statistical modelling of isotope composition is not dependent on a precise knowledge of physical or chemical laws. Instead, it is driven by linear approximation of the unknown relationships between proxies that are easy to measure and the observed isotope composition. Two interconnected questions thus emerge: which predictors should we consider (latitude, longitude, elevation, rainfall, temperature…) and under which form should we consider them (identity, squared, logarithm…)? There is no universal answer to these questions and several model structures have been proposed in the literature (see table 1 in Bowen, 2010). Importantly, the answers are likely to depend on the amount and nature of the data used to fit the models. For example, Terzer et al. (2013) suggested that different model structures should be used to model the isotope composition of precipitation water for different world regions.

The general solution employed to identify the appropriate predictors and choose their forms when building statistical models is to constitute a small set of candidate models with varying predictors and/or form for these predictors. Then, these models are fitted and their predictive power compared.

Users may decide to perform a model selection based on this model comparison. A common practice is to select the model which is best according to some criterion. This practice is controversial because when combined with null hypothesis testing it leads to an increase in the rate of false positives beyond the nominal level (Claeskens and Hjort, 2008, chapter 7 section 4; Mundry and Nunn, 2008). That being said, if the goal is not to identify whether a given predictor has a significant impact on the outcome or not, but merely to obtain a model with high predictive power, selecting the model with the highest predictive power is legitimate. An interesting alternative to model selection is to opt for a multi-model inference in which different models are retained but contribute to future predictions to a degree dependent on how good each model is. However, it is not always clear how to weight the models (Claeskens, 2016; Claeskens and Hjort, 2008, chapter 7).

The R-squared statistic has often been used to compare models. However, this is never a good measure of the ability of a model to predict new observations, even for simple linear models (*e.g.*, Faraway, 2014, p. 127). Alternative methods to estimate such predictive power have been developed for model selection, including cross-validation and the computation of various information criteria such as the Akaike Information Criterion (AIC). We now focus on two such methods, suitable in a mixed model context: the leave-one-out cross-validation (Shao, 1993) and the computation of the conditional Akaike Information Criterion (cAIC) (Vaida and Blanchard, 2005).

#### 2.5.1 Leave-one-out cross-validation

A general method to assess the predictive power of any model is to perform a cross-validation procedure where the model is used to predict data that have been observed but that have not been considered for fitting the model. When a single dataset is available, it is possible to apply a so-called leave-one-out cross-validation. In this case, one observation is excluded from the dataset (called out-of-bag observation) before fitting the model, then a prediction is generated by taking the predictors values attached to the out-of-bag observation, and this prediction is finally compared to the actual observation. This last operation is repeated by considering in turn all observations as out-of-bag observations and a metric is used to measure how close out-of-bag predictions are from their corresponding actual values (*e.g.* the mean squared error). This solution is very general and has already been used to compare other statistical approaches used to fit isoscapes ^4^

#### 2.5.2 The cAIC

The fact that our methodology is based on mixed models offers an appealing alternative to compare different models. Indeed, likelihood-based metrics called information criteria exist for such models. Information criteria have been specifically developed to allow for the comparison of the predictive power of alternative candidate models. The higher the predictive power of a given model is, the smaller the numerical values of the information criterion should be. Compared to any cross-validation procedure, the computation of an information criterion does not require to refit the models and is thus not computationally intensive.

The most famous information criterion is the AIC, which can be defined as an estimator the Kullback-Leibler divergence, *i.e.* a measure of distance between two probability distributions, the fitted model and the true unobserved reality (*i.e.*, the distribution of new observations to be predicted) (Akaike, 1973).

In the context of isoscapes, the relevant metric to compute is the *conditional* AIC (cAIC) (Vaida and Blanchard, 2005). It is so-called because it measures the predictive power of new observations drawn conditionally to the realization of the random effects that subtends the data, while the original AIC (also called *marginal* AIC) would measure the power of the fitted model to predict new response values drawn for new realizations of the random effects. Importantly, the cAIC asymptotically selects the same model than the one that selected by leave-one-out cross-validation in the context of LMM (Fang, 2011).

#### 2.5.3 An example for model comparison

As an illustration, we use our data from January (*i.e.* `GNIPdataEU1`) and compare the ability for several pairs of fitted mean and residual dispersion models to predict accurately the observed mean isotope values. We will do so using both the leave-one-out cross-validation and the cAIC. We consider three possible fixed-effect structures for the mean model and for the residual dispersion model. Specifically, we consider a fixed-effect structure defined by i) an intercept only; ii) an intercept, a linear effect of the latitude and a linear effect of the elevation; or iii) an intercept, both a linear and a quadratic effect of the latitude and a linear effect of the elevation. For the intercept-only models, we also consider two alternative random-effect structures. The first random-effect structure consists in omitting all random effects, and the second is the one introduced above (*i.e.* with the spatial and the uncorrelated random effects). For the models with more complex fixed-effect structures, we always consider the two random effects. Using these four possible model structures for each model, we will fit all 16 possible pairs of models and compare their predictive power.

For each pair of models, we proceed to the measurement of the predictive power by leave-one-out cross-validation as follows: i) we exclude a monitoring station; ii) we fit the given pair of models (*i.e.* we fit the residual dispersion model, predict the residual dispersion variance, and fit the mean model); iii) we predict the mean isotope value at the out-of-bag location; iv) we compare this value to the corresponding empirical estimate; v) we extract the squared difference, as well as the absolute difference, between these values. We repeat these five steps for each station. When this is done, we then compute the root of the average of the squared differences we obtained, as well as the average of the absolute differences, over all leave-one-out samples. We call these two averages the Root Mean Squared Error (RMSE) and the Mean Absolute Error (MAE), respectively. The smaller these error terms are, the better the predictive power.

To compute the information criterion, we fit the 16 pairs of models aforementioned to the entire dataset `GNIPdataEU1`. We then use the cAIC estimate provided by the function AIC from the package `spaMM` for the mean fit. Because this cAIC value is computed given the predictions of the residual dispersion fit, we need to penalized it by adding twice the number of parameters of the latter fit to the initial cAIC value (see eq. 6 in Overholser and Xu, 2014). The number of parameters is here computed as the sum of the number of parameters estimated for the fixed effects, plus the number of variances estimated for the random effects, plus the number of parameters estimated for the Matérn correlation function. The cAIC thereby corrected provides a measure of the joint fit to predict the response of the mean fit model.

### 2.6 Comparison with the BWR method

Many applications rely on isoscapes produced by the fitting procedure introduced by Bowen and Wilkinson (2002) and refined by Bowen and Revenaugh (2003). We call this procedure the BWR method. The BWR method is similar to ours in that it contains fixed effects accounting for the influence of covariates upon the response, but it differs from our approach in three important ways.

First, instead of estimating this spatial structure jointly with fixed effects as it is done during the fit of a mixed model, the authors proposed to fit the spatial structure on the residuals of the fit of the linear model they used to estimate fixed effects. This bears no simple relationship to likelihood estimates obtained by a joint estimation of fixed and random effects, as is apparent from the following differences.

Second, the estimation of the fixed effects is not dependent on the estimates of the random-effect parameters in the BWR method, and thus estimates are not updated iteratively during the fitting procedure, in contrast to mixed model fits. Cressie (1993, p. 22) showed that, although the estimates of the fixed-effect parameters obtained in this way remain unbiased, the standard error of these parameters are biased downwards. Because of this, tests and confidence intervals are invalid and this potentially affects prediction variance in unknown ways. A partial patch to this problem, not widely used in the isoscape literature, is to refit the fixed effects using weights derived from the spatial model (*e.g.* Zimmerman and Stein, 2010). In contrast to this patch, the full-likelihood approach we use performs such refitting until convergence of the joint estimates.

Third, a fundamental difference stems from how the spatial structure is modeled. The BWR method approximate the spatial structure by an interpolation method based on an inverse distance weighting. More precisely, in Bowen and Revenaugh (2003), the spatial component of the prediction in some given location *g*′ is computed from the vector **r** of the observed residuals *r*_*g*_ in all fitted locations, according to

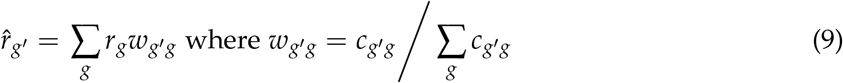

where the sum computed is other all original locations, 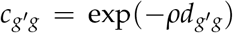 where *d*_*g′g*_ is geographical distance between the focal location *g*′ and a given location *g*, and *ρ* is a scale parameter (denoted 1/*β* in the original paper). *ρ* was estimated jointly with the regression parameters by minimizing the mean squared distance between isoscape response and isoscape predictions in the original locations. In contrast, in the mixed model the prediction of the random effect in some location *g*′ is (*e.g.* Cressie, 1993, p. 100)

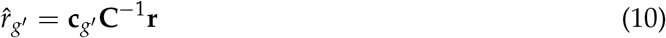

where *c*_*g*′_ is a vector of (Matérn) correlations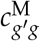, of the spatial random effect, between the focal position *g*′ and the original positions, and C is the full covariance matrix of the random part of the model, including both the random effects (spatial or uncorrelated) and the residual error. Each 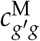 is a correlation given by the Matérn model, which becomes identical to the above exponential term *c*_*g′g*_ if the Matérn smoothness parameter is 0.5. However, the matrix C^-1^, which does not depend on new locations *x*, cannot be equivalent to the scalar 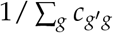. As a concrete implication of this difference, the mixed-model 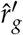 may be higher or lower than all the *r*_*g*_ values (“non-convex” prediction), which is not the case for prediction according to eq. 9 (because all *W*_*g′*g_s are between 0 and 1).

There is no definitive argument proving that the mixed-model procedure will always be better than the BWR approach. The mixed-model prediction according to eq. 10 is the best (*i.e.*, minimum-variance and unbiased) linear prediction under the assumptions of the underlying probability model embodied in the concept of a Gaussian random field, but the random field model may not be suitable in all applications. However, in most applications, the maximum of the response surface must be above observed values, so a non-convex prediction method seems appropriate. That mixed models explicitly account for residual error during the smoothing procedure should also be an advantage.

To compare the methods in practice, we implemented the BWR method as described in Bowen and Revenaugh (2003) and measured its predictive power on `GNIPdataEUl`. Here, only the leave-one-out analysis can be applied as the absence of likelihood from the BWR method prevents the computation of an information criterion.

## 3 RESULTS

### 3.1 The fitted models

To illustrate how to model the spatial distribution in isotope composition using mixed-models, we have fitted the mean model `meanfit1` and the residual dispersion model `dispfit1` on a aggregated dataset containing the estimated mean and inter-annual variance in *δ*^2^H found in precipitation water across Europe for the month of January (`GNIPdataEU1`). In this first section, we explain how to interpret the estimates obtained for such models.

#### 3.1.1 The mean model

The mean model we chose only contains three terms to model the fixed effect `Fix` – the intercept, the slope for the effect of latitude and the slope for the effect of the elevation – which were estimated as 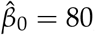, 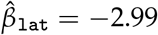 and 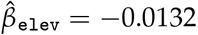, respectively (the estimates for all twelve months are provided in table 4). The intercept represents the point prediction in unit of *δ*^2^H for a zero latitude, zero elevation and a realization of random effects equal to zero. As such, it is not easily related to an observed feature of the predicted isoscape. In contrast, the slope estimates represent trends that can be perceived from the fitted isoscape, although they are blurred in any location by the random effects. Precisely, the estimates 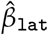 and 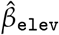 predict the change in the mean isotope composition associated with an increase in one degree of latitude and an increase in one meter of elevation, respectively. For example, an increase in elevation of 1000 meters is predicted to shift the mean isotope value by −13.2 *δ*^2^H.

**Table 4.** Sample sizes and parameter estimates for all 12 fitted mean models. The columns N_o_b_s_ and N_station_ provide the total number of observations used in each model and the total number of measuring stations, respectively. Different observations per station always correspond to different years. The meaning of the other columns is discussed in the text. The last two rows provide the direct mean and standard deviation for each parameter estimate.

| Model | Month | N<sub>obs</sub> | N<sub>station</sub> | $\hat{\beta}_0$ | $\hat{\beta}_{\text{lat}}$ | $\hat{\beta}_{\text{elev}}$ | $\hat{\lambda}_{\text{RS}}$ | $\hat{\lambda}_{\text{RU}}$ | $\hat{\rho}$ | $\hat{v}$ |
| --- | --- | --- | --- | --- | --- | --- | --- | --- | --- | --- |
| meanfit1 | January | 2759 | 313 | 79.99 | -2.99 | -0.0132 | 5415.5 | 9.02e-11 | 2.26e-05 | 0.384 |
| meanfit2 | February | 2697 | 308 | 84.15 | -2.88 | -0.0154 | 1547.9 | 6.67e-11 | 6.60e-05 | 0.306 |
| meanfit3 | March | 2702 | 308 | 85.89 | -2.85 | -0.0150 | 1088.1 | 9.31e-11 | 9.54e-05 | 0.346 |
| meanfit4 | April | 2583 | 291 | 83.17 | -2.53 | -0.0141 | 552.5 | 1.12e-10 | 1.00e-04 | 0.322 |
| meanfit5 | May | 2494 | 272 | 66.82 | -2.12 | -0.0126 | 220.1 | 1.70e-10 | 2.08e-04 | 0.371 |
| meanfit6 | June | 2322 | 236 | 65.54 | -2.07 | -0.0088 | 102.8 | 4.80e+00 | 4.73e-04 | 0.500 |
| meanfit7 | July | 2203 | 225 | 65.55 | -1.98 | -0.0068 | 107.0 | 2.25e-10 | 4.58e-04 | 0.372 |
| meanfit8 | August | 2226 | 238 | 78.54 | -2.29 | -0.0077 | 143.3 | 2.71e-10 | 4.06e-04 | 0.471 |
| meanfit9 | September | 2453 | 251 | 70.81 | -2.28 | -0.0109 | 221.7 | 1.99e+00 | 4.10e-04 | 0.491 |
| meanfit10 | October | 2635 | 265 | 83.26 | -2.67 | -0.0117 | 530.7 | 1.78e-10 | 2.44e-04 | 0.498 |
| meanfit11 | November | 2692 | 293 | 79.24 | -2.85 | -0.0134 | 2092.8 | 8.66e-11 | 3.39e-05 | 0.333 |
| meanfit12 | December | 2780 | 313 | 70.25 | -2.78 | -0.0132 | 4197.0 | 1.21e-04 | 1.48e-05 | 0.324 |
| MEAN |  |  |  | 76.10 | -2.52 | -0.0119 | 1351.6 | 5.67e-01 | 2.11e-04 | 0.393 |
| SD |  |  |  | 7.44 | 0.34 | 0.0027 | 1676.0 | 1.39e+00 | 1.73e-04 | 0.072 |

The variances for the two random effects of the mean model are estimated to 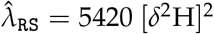 and 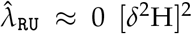, for `RandomSpatial` and `RandomUncorr`, respectively. The estimate 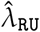 is negligible, which suggests that there is no particular bias in the measurements introduced by the measuring stations. The different water samples collected at a given station can thus be considered as independent once the effect of the location of the station is accounted for. In contrast, the value of 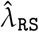 is not negligible. The spatial covariances also depend on the spatial correlation parameters, and the best way to study spatial covariation in an isoscape is thus to directly visualise the predictions of the random effects as we will do below.

#### 3.1.2 The residual dispersion model

The residual dispersion model we chose only contains a single term to model the fixed effect `Fix`^D^ - the intercept 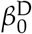 - which was fitted to 5.26 (the estimates for all twelve models are provided in table 5). As explained above, the intercept is not particularly meaningful in the present context.

**Table 5.** Sample sizes and parameter estimates for all 12 fitted residual dispersion models. See table 4 for legend. Note that the sample sizes presented here are slightly lower than those of the mean models (table 4). This is because the month-location combinations only available for a single year do not yield to variance estimates `variso` and are thus not used while fitting the residual dispersion models. Once the residual dispersion models are fitted, it is however possible to predict the residual dispersion for these particular combinations, which is why they are being considered during the fit of the mean models.

| Model | Month | N<sub>obs</sub> | N<sub>station</sub> | $\hat{\beta}_0^D$ | $\hat{\lambda}_{RS}^D$ | $\hat{\lambda}_{RU}^D$ | $\hat{\rho}^D$ | $\hat{v}^D$ |
| --- | --- | --- | --- | --- | --- | --- | --- | --- |
| dispfit1 | January | 2696 | 250 | 5.26 | 4.61 | 1.00e-06 | 1.85e-05 | 0.328 |
| dispfit2 | February | 2631 | 242 | 5.46 | 0.47 | 1.00e-06 | 4.01e-04 | 0.309 |
| dispfit3 | March | 2627 | 233 | 5.47 | 0.88 | 9.99e-04 | 4.11e-05 | 0.236 |
| dispfit4 | April | 2517 | 225 | 5.49 | 0.77 | 1.00e-06 | 1.89e-05 | 0.163 |
| dispfit5 | May | 2426 | 204 | 5.73 | 5.30 | 1.00e-06 | 1.48e-05 | 0.399 |
| dispfit6 | June | 2287 | 201 | 5.62 | 7.49 | 1.00e-06 | 1.99e-05 | 0.488 |
| dispfit7 | July | 2163 | 185 | 5.34 | 3.30 | 1.00e-06 | 1.48e-05 | 0.343 |
| dispfit8 | August | 2184 | 196 | 5.04 | 0.21 | 1.00e-06 | 3.07e-03 | 0.500 |
| dispfit9 | September | 2409 | 207 | 5.16 | 2.78 | 1.00e-06 | 1.48e-05 | 0.322 |
| dispfit10 | October | 2600 | 230 | 5.21 | 0.96 | 1.00e-06 | 9.88e-05 | 0.288 |
| dispfit11 | November | 2637 | 238 | 5.47 | 2.45 | 1.00e-06 | 1.49e-05 | 0.300 |
| dispfit12 | December | 2719 | 252 | 5.46 | 0.53 | 3.09e-02 | 2.72e-04 | 0.500 |
| MEAN |  |  |  | 5.39 | 2.48 | 2.66e-03 | 3.33e-04 | 0.348 |
| SD |  |  |  | 0.19 | 2.22 | 8.51e-03 | 8.33e-04 | 0.101 |

The variances for the two random effects of the residual dispersion model are estimated to 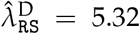 and 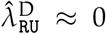, for, `RandomSpatial`^D^ and `RandomUncorr`^D^, respectively^5^ As for the mean model, the estimate of the uncorrelated random term 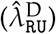 is negligible and the estimate of the spatial random term 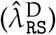 should be investigated jointly with the estimates of the Matérn parameters by simple visualisation of the predictions of the random effects.

### 3.2 Isoscapes

In fig. 1, we show three isoscapes that can be built from mixed model fits, which are relevant for interpreting the spatial variation in isotope composition. The coarse resolution of the maps produced here is only stemming from the coarse elevation raster we used and it is possible to use high resolution elevation rasters to produce very detailed isoscapes. In this figure, the isoscapes associated with the fitted models for January (*i.e.* based on `meanfit1` and `dispfit1`, see table 4 & 5) are shown in the first row of fig. 1a-c. The second row provides corresponding isoscapes for the month of July (*i.e.* based on `meanfit7` and `dispfit7`) and thus illustrates seasonal variation. The last row of fig. 1 shows the three isoscapes for the precipitation-amount weighted annual averages which we built using precipitation data from WorldClim 1.4 (Hijmans et al., 2005). We will limit our description to the first three isoscapes, as other can be interpreted similarly.

The first isoscape (fig. 1a) represents the point predictions for *δ*^2^H in Europe for the month of January. The isoscape of point predictions is usually the one of main interest as it describes how the isotope composition varies in space. Here, the patterns of the *δ*^2^H value looks similar to what has been documented by others when using alternative approaches: *δ*^2^H values decrease along the axis SW/NE and they also decrease with elevation. The second isoscape (fig. 1b) represents the prediction variances associated to the point predictions. This isoscape shows where point predictions are most unreliable. Here, North Africa and the South-East corner of the studied geographic area are associated to high prediction variances, which is not surprising as these regions are disjunct from the rest by large bodies of water which impact on the isotope composition and they are simultaneously associated with few observations. The third isoscape (fig. 1c) represents the residual variances associated to the first isoscape. This isoscape shows where *δ*^2^H values varies the most between year. Here, we see that more inter-annual variation in *δ*^2^H values occurs in Russia and around the Caspian sea during January.

**Fig. 1.**
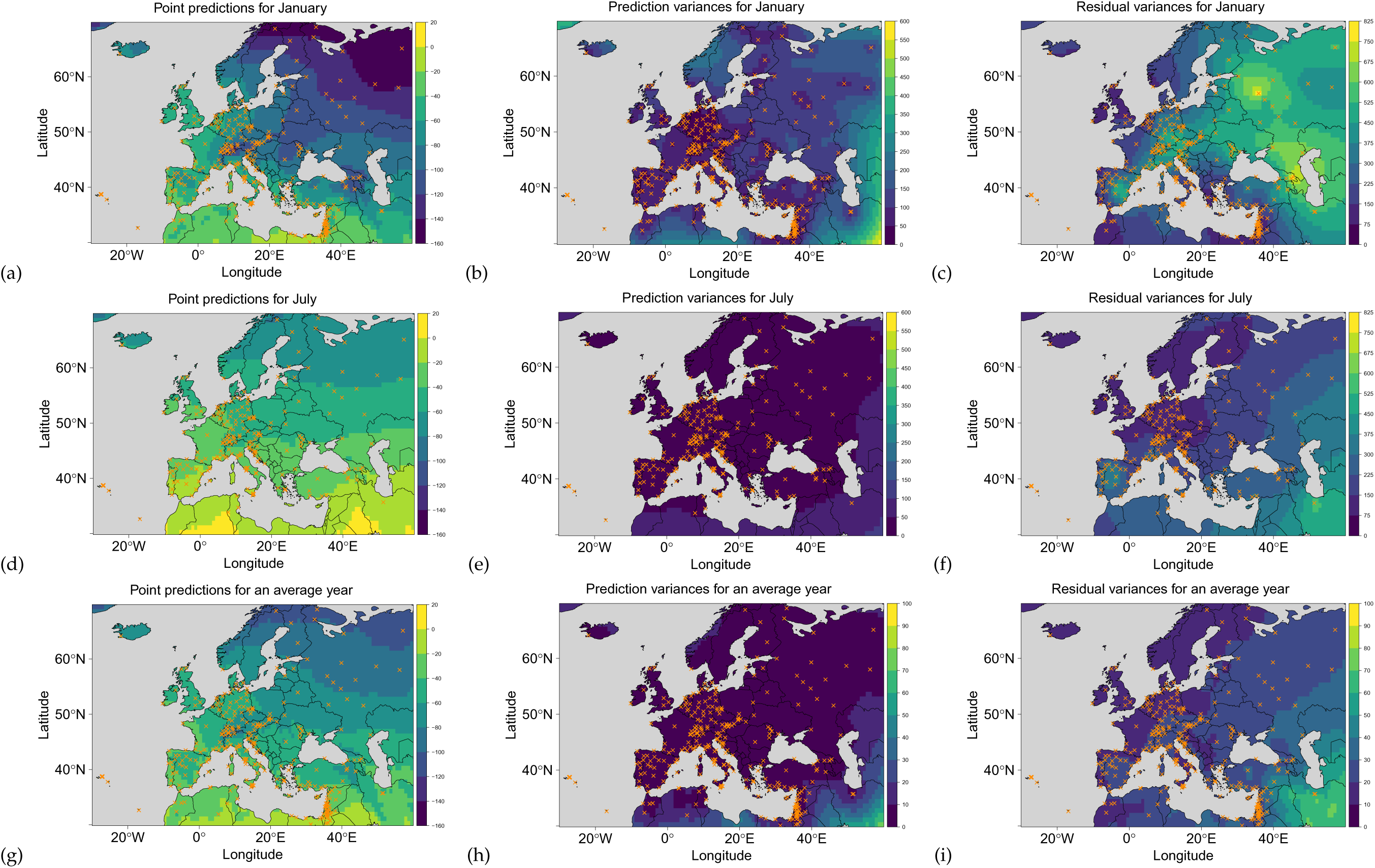
Isoscapes for the isotope composition of the hydrogen from precipitation water during the period 1960-2014. The first column provides point predictions (a, d, g), the second column provides prediction variances (b, e, h) and the third column provides residual variances (c, f, i). These isoscapes depict this information for the *δ*^2^H values collected during January (a, b, c), July (d, e, f), as well as for the predpitation-amount weighted annual averages (g, h, i). Colours depict *δ*^2^H values (a, d, g) or [*δ*^2^H]^2^ values (b, c, e, f, h, i). Note that the colour scales differ between some of the plots.

#### 3.2.1 Isoscapes for the fixed and random effects separately

The previous isoscapes represent joint predictions stemming from both fixed and random effects. It is however possible to visualise predictions for the fixed and for the random effects separately. We illustrate this possibility for the mean model in fig. 2. The sum of these two isoscapes equates the isoscape for point predictions shown in fig. 1a. Visualising the fixed effects predictions in isolation reveals the geographical patterns captured by predictors even in the presence of strong spatial autocorrelation. Here, the isoscape for the mean model confirms what we discussed above: the *δ*^2^H values decrease with latitude and elevation (fig. 2a). In turns, visualising the mean of the conditional realizations for the random effects reveals the geographical patterns of the spatial autocorrelation. Here, it shows that spatial autocorrelation spans over large distances (fig. 2b), resulting in an increase in *δ*^2^H values close to the Atlantic ocean (except in Africa) and a decrease in values when the distance to the Atlantic increases. We find this latter isoscape particularly useful as no numerical output from our fitted models would have allowed us to reach this conclusion efficiently. In contrast, the visualisation of the fixed effects in isolation merely offers a simple alternative to the interpretation of the numerical values of the fixed effect estimates. Importantly, while we can represent separately the contribution to point predictions from fixed effects and random effects, we recall that the fit of the mean model is a *joint* fit of the fixed and random effects: both components were thus fitted simultaneously and not separately.

**Fig. 2.**
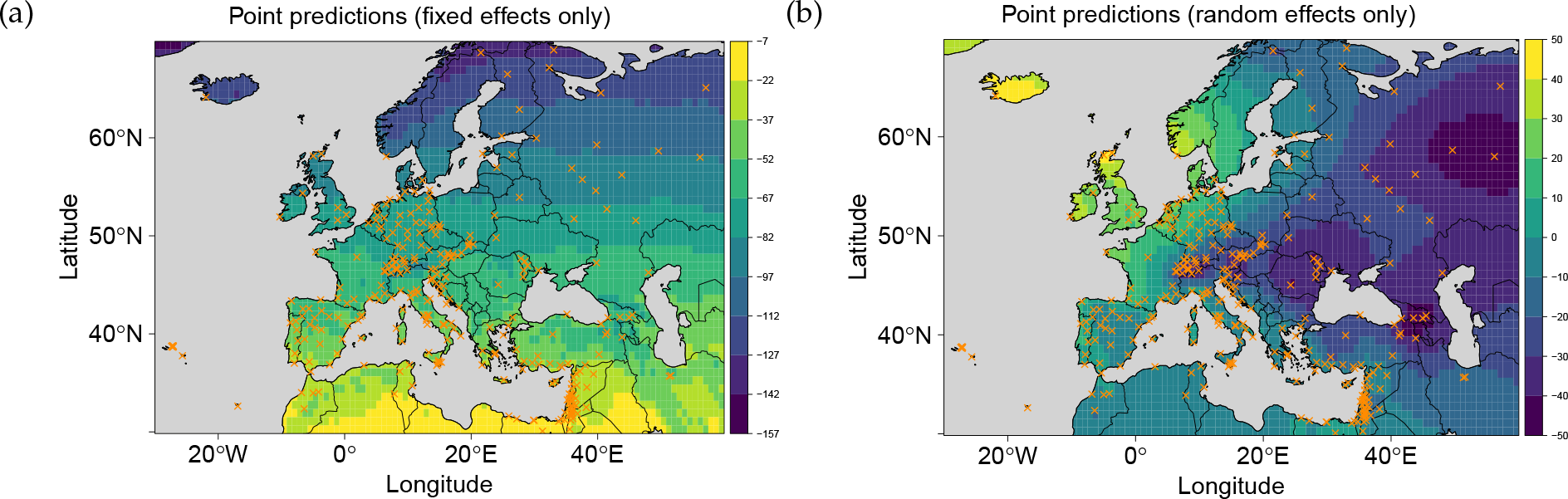
Decomposition of the predictions for `meanfit1` in terms of fixed (a) and random effects (b). The sum of these two maps gives the isoscape for point predictions shown in fig. 1a. The colours depict *δ*^2^H values and oranges crosses depict sampling locations.

### 3.3 Model comparison and model selection

We compared the predictive power associated with the fits of 16 different pairs of mean and residual dispersion models (table 6). To assess the predictive power we used three different metrics: the root mean squared error (RMSE) and the mean absolute error (MAE) which we computed by leave-one-out cross-validation, and the cAIC which is analytically derived from single fits. Two main results emerged. First, results show that considering random effects largely improves the predictive power of the fits. Indeed, all fits considering a mean model with random effects show a predictive power higher than the those associated with a mean model without any random effects. For the fits in which the mean model considered random effects (*i.e.* B, C & D; see table 6 for notation), the predictive power is similarly improved when these models have been fitted given a residual dispersion model also considering random effects. The estimates of the variance of the uncorrelated random effect are generally low. Therefore, the gain in predictive power triggered by the random effects results from the spatially autocorrelated random effect.

**Table 6.** Model comparison between four different fixed-effect parametrizations used for the fit of the mean model and the residual dispersion model on the January data. The first column provides a two-letters code which refers to the formula used for fitting the mean model (upper case) and the residual dispersion model (lower case). For the mean model, we consider the following formulas: A = *β*_0_, B = *β*_0_ + *R*, C = *β*_0_ + *β*_lat_ × `lat`_*g*_ + *β*_elev_ × `eleV`_*g*_ + *R*, and 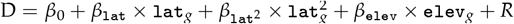. For the residual dispersion models, we similarly considered: 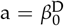, 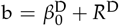, 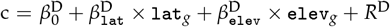, and 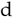 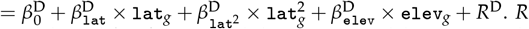 and R^D^ refer to the spatial and uncorrelated random terms for the mean models and for the residual dispersion models, respectively. We also included the model fitted according to Bowen and Revenaugh (2003) for comparison and call this model BWR. The second and third columns give the number of parameters estimated in the mean model (K_mean_) and in the residual dispersion model (K_disp_). The fourth column gives the conditional Akaike Information Criterion (cAIC) before correcting it by K_disp_. The fifth column gives such corrected cAIC value. The sixth and seventh columns give the Root Mean Squared Errors (RMSE) and the Mean Absolute Error (MAE), respectively. These two measurements of the prediction errors were computed by leave-one-out cross-validation. The last three columns give the model ranks according to each metric used for model comparison (the smaller the rank, the better the predictive power). Note that ranks may differ between RMSE and MAE although values for these metrics appear identical because the ranks are influenced by further decimal places than those displayed. We do not provide information criterion for the BWR model due to the absence of likelihood associated to this method.

| models | $K_{\text{mean}}$ | $K_{\text{disp}}$ | $\text{cAIC}_{\text{uncorrected}}$ | cAIC | RMSE | MAE | $\text{rank}_{\text{cAIC}}$ | $\text{rank}_{\text{RMSE}}$ | $\text{rank}_{\text{MAE}}$ |
| --- | --- | --- | --- | --- | --- | --- | --- | --- | --- |
| Aa | 1 | 1 | 9287.00 | 9289.00 | 32.99 | 28.10 | 13 | 14 | 17 |
| Ab | 1 | 5 | 11584.52 | 11594.52 | 33.57 | 26.80 | 14 | 15 | 14 |
| Ac | 1 | 7 | 11597.63 | 11611.63 | 33.59 | 26.81 | 15 | 16 | 15 |
| Ad | 1 | 8 | 11616.67 | 11632.67 | 33.59 | 26.81 | 16 | 17 | 16 |
| Ba | 5 | 1 | 2365.43 | 2367.43 | 13.71 | 9.49 | 12 | 13 | 13 |
| Bb | 5 | 5 | 2292.65 | 2302.65 | 13.69 | 9.48 | 7 | 10 | 11 |
| Bc | 5 | 7 | 2291.89 | 2305.89 | 13.69 | 9.48 | 8 | 11 | 10 |
| Bd | 5 | 8 | 2292.30 | 2308.30 | 13.70 | 9.49 | 9 | 12 | 12 |
| Ca | 7 | 1 | 2346.76 | 2348.76 | 12.76 | 8.78 | 10 | 7 | 7 |
| Cb | 7 | 5 | 2273.36 | 2283.36 | 12.70 | 8.68 | 1 | 1 | 2 |
| Cc | 7 | 7 | 2272.93 | 2286.93 | 12.70 | 8.68 | 3 | 2 | 1 |
| Cd | 7 | 8 | 2273.28 | 2289.28 | 12.70 | 8.68 | 5 | 3 | 3 |
| Da | 8 | 1 | 2346.93 | 2348.93 | 12.79 | 8.79 | 11 | 8 | 8 |
| Db | 8 | 5 | 2273.41 | 2283.41 | 12.72 | 8.69 | 2 | 4 | 5 |
| Dc | 8 | 7 | 2272.98 | 2286.98 | 12.72 | 8.69 | 4 | 5 | 4 |
| Dd | 8 | 8 | 2273.33 | 2289.33 | 12.73 | 8.69 | 6 | 6 | 6 |
| BWR | 5 | 0 |  |  | 13.05 | 9.20 |  | 9 | 9 |

Second, within the subset of pairs of models considering random effects (*i.e.* Bb, Bc, Bd, Cb, Cb, Cd, Db, Dc, Dd), the fits involving environmental predictors for the mean model (models C or D) show the highest predictive power. In sum, our model comparison shows that the best fits of isotope values are obtained when we consider both environmental predictors as fixed effects and spatial autocorrelation as random effects.

Among the mean models considering environmental predictors, all metrics (*i.e.* RMSE, MAE & cAIC) identify the mean model C as best. The exact ranking of the predictive power for the pairs of fits involving the mean model C differs between MAE and the two other metrics. However, the two best models (Cb and Cc) have practically identical RMSE and MAE (see table 6) and cAIC then retains the most parsimonious of these two models. The pair Cb is the one we presented in the introduction.

That all metrics support the same general findings and that they all lead to select the same model suggests that one could solely rely on cAIC to select the best models. This would allow one to avoid the computational cost of a cross-validation which is particularly large in the context of mixed models. Here for example, we obtained cAIC values in around an hour for each fit, but the computation of the RMSE and the MAE required several thousands CPU hours. The empirical equivalence we found between the cAIC and the cross-validation is expected to hold asymptotically for LMMs (Fang, 2011), but remains to be formally established for GLMMs.

### 3.4 Comparison with the BWR method

We now compare the prediction errors measured during the leave-one-out cross-validation between our mixed-models based approach and the BWR method (fig. 3 & 4). Table 6 shows that the BWR method leads to lower prediction accuracy than any pair of mixed models accounting for environmental covariates (Ca, Cb, Cc, Cd, Da, Db, Dc, Dd). For either approaches, there is no obvious spatial structure in the distribution of the errors (fig. 4a & 4b). Yet, the prediction errors between the two approaches appear highly correlated (fig. 3c), which means that both approaches tend to produce overestimate (or underestimate) the *δ*^2^H values at the same measuring stations.

**Fig. 3.**
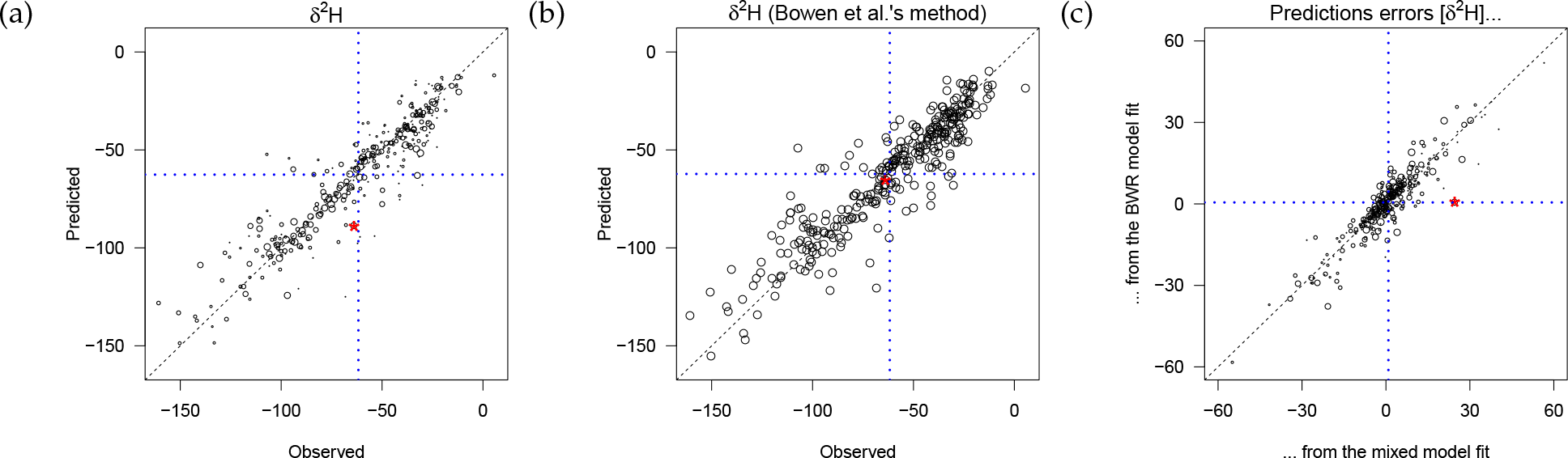
Assessment of predictive power by leave-one-out cross-validation for January. The first two plots compare predictions and observations for out-of-bag observations when predictions have been generated by the best fits Cb selected in table 6 (a), and by the BWR fit (b). The last plot (c) compares prediction errors between our best fits and the BWR fit. In the plots produced by mixed models (a & c), the diameter of the circles is proportional to the logarithm of the prior weight used during the fit. The outlier corresponding to a sampling location in Iceland (see text for details) is represented by an asterisk. The black dashed line represents the identity relationship, so points falling on this line correspond to locations where the predicted and the observed values are identical (a & b) or to locations where the prediction error is the same for our approach and for the one by Bowen and colleagues (c). The horizontal and vertical blues lines represent the raw mean of the data along each axis.

**Fig. 4.**
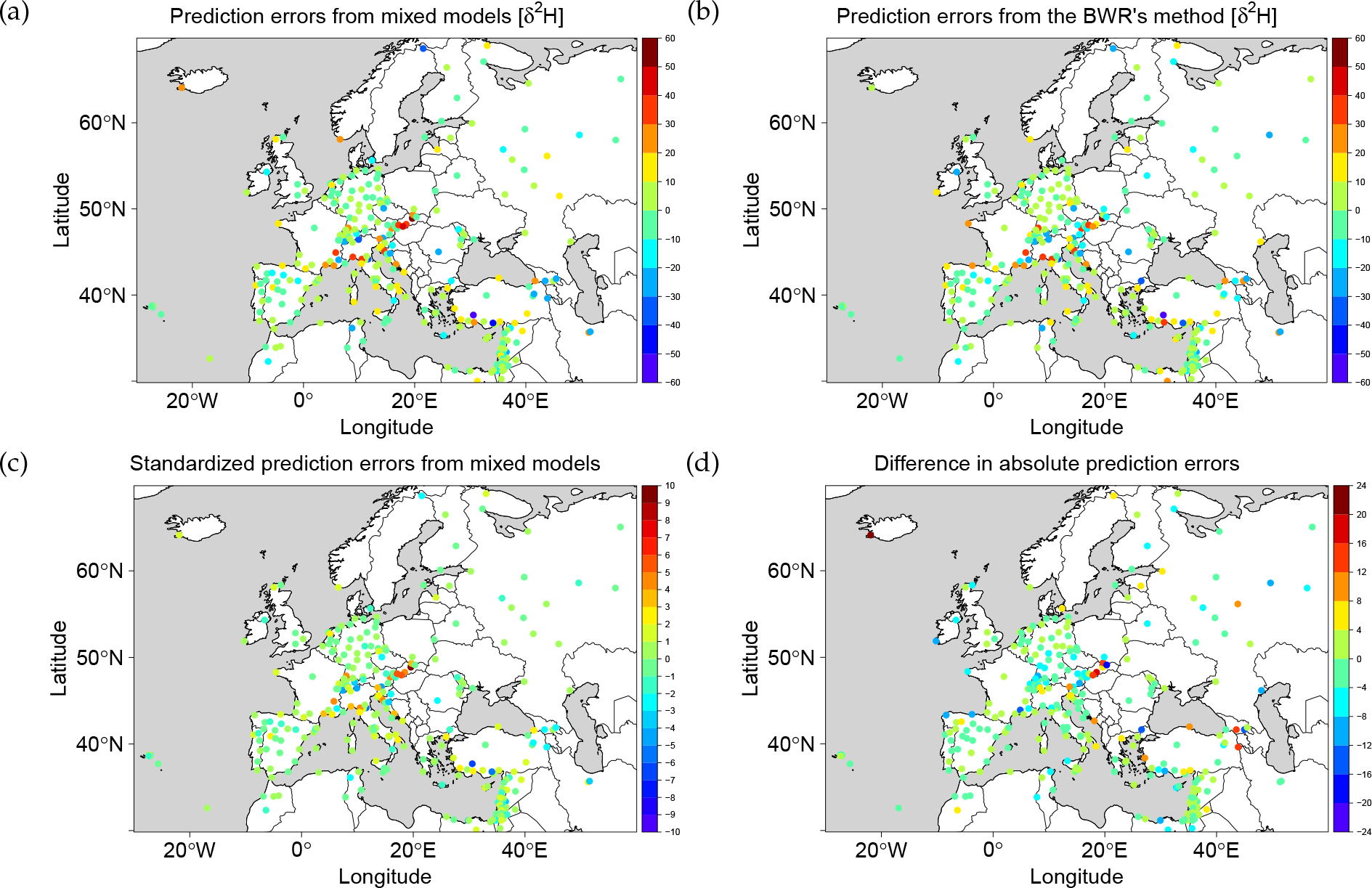
Spatial distribution of the prediction errors during the leave-one-out cross-validation for January. We show the prediction errors (*i.e.* observed-predicted value at the out-of-bag observation) for the mixed models based approach (a) and for the approach followed by Bowen and colleagues (b). The plot (c) represents the same data as plot (a) but here the predictions errors have been divided by the square root of the prediction variance computed at the corresponding out-of-bag locations. The last plot (d) represents the differences in absolute prediction errors between the mixed models and those from the approach followed by Bowen and colleagues.

One notable exception is the point represented by an asterisk in fig. 3c which seems much better predicted by the BWR method than by mixed models. This outlier illustrates a fundamental difference between the two approaches. The point corresponds to measurement from Iceland, which is a location distant from all other observations (fig. 4d). During the leave-one-out crossvalidation the models are refitted without this point and thus the prediction associated to this point represents an extrapolation. Here, the observation turns out to have a *δ*^2^H value close to the mean of all the over observation (fig. 3), but due to spatial autocorrelation our method predict a large positive value for the realization of the random effect in this location (fig. 2b). In contrast, the BWR method predicts mean values at location far from observations. As a consequence, the BWR method turns out to predict this out-of-bag observation very well and our approach does not. Had the observed value for Iceland been quite different from the mean of all other observations, the BWR would have made a large prediction error for this observation too.

In contrast to the BWR method, our approach quantifies the uncertainty associated with predictions. In fact, the prediction variance at the location of the measuring station in Iceland (in the fit considering this location as out-of-bag) is 487.3 [*δ*^2^H]^2^, which is the highest prediction variances we obtained during the cross-validation (it is higher than the second largest prediction variance by 158.7 [*δ*^2^H]^2^). This illustrates that our method allows for the identification of locations where prediction errors are likely to happen. Once the prediction variance is accounted for, the prediction errors do not seem to present any strong spatial structure (compare fig. 4a and 4c).

## 4 DISCUSSION

In this paper we explained how the spatial distribution of the isotope composition can be fitted using mixed models. We have illustrated this by constructing isoscapes of the isotope composition of hydrogen (*δ*^2^H) for precipitation water in Europe by the mean of a general mixed model `R` package (`spaMM`: Rousset and Ferdy, 2014). We will now discuss the potential benefits and constraints associated to this novel approach.

### 4.1 Mixed model vs the BWR method

We have compared our mixed model approach to one of the most widely used approach in the field: the BWR method (Bowen and Revenaugh, 2003; Bowen and Wilkinson, 2002). We have detailed theoretical reasons for the mixed model approach to be better. Accordingly, we found that, at least on the dataset we used, the mixed models approach predicts better unobserved isotope measurements.

The mixed model approach is much slower to fit than the BWR fitting procedure. In our example, the BWR fitting procedure was up to 3 orders of magnitude faster than the mixed model approach. This is however not a major practical limitation since the aggregation scheme we propose still allows for a complete fit in a few hours maximum on a personal computer. Most users are likely to rely on dataset smaller than the one we used, bringing the time cost down. In addition, although the fitting procedure is slow, because mixed models provide a measure of the likelihood, a single fit is necessary to assess the prediction accuracy of such a fit. We have indeed shown that an information criterion such as the cAIC could be used for such purpose. In contrast, the prediction accuracy of a BWR fit must be assessed by cross-validation which requires to refit the model multiple times.

Another small caveat of a mixed-model based approach is that it may poorly extrapolate, as we have seen it for the case of Iceland. However, we have shown that this problem is directly diagnosed by the quantification of the prediction errors that mixed models makes possible. We have also illustrated that mixed models allow also for a quantification of the temporal variation in isotope composition through the fit of the residual variance. We will now detail why the ability for the mixed model to produce isoscapes representing such prediction variance and residual variance is a strong argument in favour of the method.

### 4.2 The utility of isoscapes for prediction variance and residual variance

That our method produces isoscapes of both the prediction and residual variances presents several interests in terms of applications. First, the isoscape representing prediction variance is informative about the sampling design: it directly informs where predictions are associated with the highest uncertainty, and therefore where to collect data in order to reduce such uncertainty. It also reflects the overall level of uncertainty for an isoscape. Here for example, it shows that monthly isoscapes are much more inaccurate than annual isoscapes. Thus, practical implications directly follows from the examination of isoscapes representing the prediction variance. Authorities such as the Global Network of Isotopes in Precipitation (IAEA/WMO, 2017) could use them to establish priorities for the extention of their existing network. Moreover, annual isoscapes and not monthly isoscapes should probably be used for application where precision is paramount, such as infering location of migratory organisms.

Second, the isoscape representing residual variances is informative about temporal variation in isotope composition. In our study, the response variable of the residual dispersion model (`variso`) corresponds to inter-annual variation in *δ*^2^H and thus the isoscape of the residual variance shows this variation. Modifying the response variable would thus allow for the study of variation in *δ*^2^H (or other isotopes) at other time scales. Our study revealed that the inter-annual variation is strongly spatially structured. We captured such spatial structure of the temporal variation by means of random effects. The consideration of suitable predictors for fixed-effects could refine further the accuracy of the fit for this spatial structure. Here, the fixed-effect parametrizations we tried did not improve noticeably the quality of the fit, compared to the mere consideration of an intercept. Other parametrizations should thus be attempted, guided by the understanding of the determinants of isotope composition and the availability of data.

Our approach also allows for the computation and display of a third kind of isoscape that directly derives from the isoscapes for prediction variance and the one for residual variance: the isoscape of the response variance. The response variance is simply the sum of the prediction variance and of the residual variance. Computing the response variance is especially interesting with respect to the inference of the origin of environmental materials based on isotope composition (Hobson and Wassenaar, 2008). For example, environmental materials originating from area with low *δ*^2^H values do not necessary come from locations where the point prediction is low. Such organisms may also come from locations where the *δ*^2^H values are only sometimes low; so from locations with a mean that is not necessarily low but with a high residual variance. They may also come from locations where point predictions are overestimated; so from locations with high prediction variance. These two possibilities are captured by the computation of the response variance. It is crucial not to neglect either sources of variance as it would lead to rule out possible origin locations, which may lead to erroneous inference, which may in turn lead to wrong decision making. That such decisions impact on areas such as conservation practices in biology or criminal investigation in forensic sciences should be a strong motivation to also study isoscapes of the response variance. We have already implemented such an approach in our package `IsoriX` (Courtiol et al., 2016) and we will explain how to perform geographical assignments accounting for both the prediction and the response variance using mixed models in a future publication.

### 4.3 Assumptions of the current approach and possible extensions

Different statistical approaches differ in the assumptions they make. It is important to make explicit these assumptions in order to anticipate in which conditions a given approach may or may not be adequate. That our approach is based on mixed models make this exercise easier than for methods that are not based on explicit assumptions. We will now discuss the assumptions that seem the most relevant in the context of isoscapes.

First of all, following a standard assumption of linear models, mixed models consider that each of the covariate values used to model fixed effects has been measured without error. This assumption is also implicitly made in alternative approaches, but it is often violated in practice because there is often measurement errors in the predictors. In particular, raster values (here, the elevation values) are often the outcome of some smoothing procedures which generate errors. The raster values do not correspond, in general, to the real value at the prediction location, as its coordinates would suggests, but to the result of a function applied on all the data measured within the cell surrounding the prediction location (often the mean). We therefore recommend for user to pay a particular attention to the source of data for their covariates and to select the rasters with the highest available resolution and most precise measurement available. Here, we used a very coarse elevation raster in order to be able to provide this raster as Supplementary material. However, our toy raster has been derived from a high resolution raster that could be used instead for real applications. This raster is the Global Multi-resolution Terrain Elevation Data 2010. It has a resolution of 7.5 arc seconds (*i.e.* 232m along the equator and 102m along the Arctic circle) and a standard deviation of the prediction error for the elevation of only 6 meters (based on comparison with control points)^6^

A second assumption is that the environmental predictors are assumed to exert a linear effect on the response. Again, this assumption also concerns alternatives (*e.g.* Bowen and Revenaugh, 2003; Bowen and Wilkinson, 2002) and transformation of the predictors can be made as long as the transformed predictors are still considered as linear in their effects. For example, the predictors can be expressed as polynomials (as we have seen for latitude) or they can be logarithmic transformed. Nonetheless, the reliance on linearity precludes the implementation of some model structures (*e.g.* the one used by Van der Veer et al., 2009).

Third, our approach also makes specific assumptions about the random effects. We have considered all random effects to be normally distributed. It is easy to consider alternative distributions (*e.g.* Gamma) for the random effects whose realizations are not spatially autocorrelated (here `RandomUncorr` & `RandomUncorr`^D^) and the package `spaMM` we used to fit the mixed models offers this possibility. However, because random effects that present autocorrelated random effects must be described in terms of mean and covariances, using alternative distributions is not currently possible for the spatial random effect (unless one considers other elliptical distributions closely related to the Gaussian; Embrechts et al., 2002). Other Kriging approaches for which assumptions have been made explicit also assume normality (Van der Veer et al., 2009).

Fourth, we also made assumptions during the computation of the prediction variance and the residual variance. For the prediction variance, the fitting procedure we used considers the estimates for the uncertainty of the estimates for fixed effects, as well as the uncertainty of the realization of the random effects. However, it does not yet allow one to account for the uncertainty stemming from the estimation of the correlation parameters and the uncertainty associated with the estimation of the variances of the random effect is only partially accounted for. Again, the same is true for other Kriging approaches which usually do not consider uncertainty in the Kriging parameters. Some bootstrap methods have been discussed in the mixed-model literature to account for these neglected sources of uncertainty (Booth and Hobert, 1998), and can be performed using `spaMM` and little additional programming, but these methods are computationally intensive.

Moreover, for the residual variance, all sources of uncertainty are being neglected as only point predictions are being used from this model in order to fit the mean model. That implies that we consider that the residual variance is exactly known. Many alternative Kriging approaches also make this assumption and we do not know of any study have relaxed such assumptions in the context of isoscapes. A considerable improvement other alternative approaches, however, is that our method do not assume such known variance to be constant. Indeed, in other works the residual variance is at worse assumed to be null and at best considered to be constant. Here, we have employed a model-based approach for the residual variance that we have modelled both using fixed effects and random effects. In particular, we have included a random effect for which realizations are spatially autocorrelated. That means that we have considered that the mean *δ*^2^H values and that the inter-annual variance in these *δ*^2^H values follow two distinct spatial processes. To compare our approach to one that would assume that the residual variance does not vary in space, we included in our models comparison a fit of the residual dispersion model considering only an intercept and no random effect (models Xa in table 6, with χ = A, B, C or D). Doing this we obtained a cAIC value considerably higher than any of our residual dispersion models considering random effects. Considering that the residual variance varies in space is thus fully justified in our context.

Fifth, we have considered that the distribution of the isotope composition is independent between between different months and also independent between different years. This is obviously not realistic as strong temporal autocorrelation may exist. Most other statistical approaches to isoscapes also neglect this temporal autocorrelation (but see Klaus et al., 2015) and further development should be made to alleviate this strong assumption. Correlation functions allowing to account for both spatial and temporal correlation exist for mixed models and could thus be used within our formalism (but this is not implemented in the current version of `spaMM`). Accounting for the autocorrelation should lead to reduction in the prediction variance and thus to more precise inference.

With this caveat in mind we showed that our approach provides information about different time scales simultaneously. In our example, one could study month-to-month variation by comparing monthly isoscapes, and year-to-year variation by looking at the isoscape of the residual variance. Nonetheless, both time scales are treated differently. While it is possible to visualize the realization of the spatial process for a given month, it is not possible to compute an isoscape corresponding to a given year. To do the latter, we do not recommend to fit a new model considering the data from a single year, as it would prevent the estimation of the residual variance which is required to compute the prediction variance. Instead, an alternative allowing for the examination of yearly isoscape would be to invert both time scales; that is, to split the data per year (instead of per month) during the aggregation process and to use as the response variable for the dispersion model the observed estimates of the inter-month variation within year for each measuring station. For such estimate to be reliable only stations from which samples are available for the same months should be retained (*e.g.* we could consider the station that have records for all months), which imply to lose some data. Here, we chose to aggregate the data accross years and not accross months because there is stronger temporal structure between months than between years.

Sixth, instead of fitting the mean and residual dispersion simultaneously, we fitted sequentially the residual dispersion model and then the mean model. We did so to allow for a considerable gain in computation time, which may be required as spatial mixed models are generally slow to fit. The gain in computation time is present for two reasons. First, our sequential procedure allows to aggregate the data and to directly work on the mean and variance estimates of each month-location combination, rather than working on the total number of water samples. Second, our procedure provides dispersion estimates which converge immediately, because the response of the dispersion model does not depend on the results of fitting the mean model for given residual variance.

Our sequential fitting approach thus differs from the full-likelihood method described by Lee and Nelder (2006, p. 327) which is iterative because therein the estimation of residual dispersion parameters depends on the results of the mean fit. They call their approach a Double Hierarchical GLM (DHGLM; Lee and Nelder, 2006). This latter approach requires to work on non-aggregated data. The package offers the possibility to fit DHGLMs, but such fits are extremely computer intensive for joint spatial models. The estimates by our sequential approach should differ from those obtained by a DHGLM. Indeed, estimates of the residual variance parameters generally differ between a sequential and a joint fitting procedure, even in the simpler case when the residual variance is fitted only by an intercept term, and Searle et al. (2006, Section 4.7) showed that differences are more generally expected for mixed models. That being said, we only expect slight differences in estimates when the parameters of the mean model and the residual dispersion models are fitted simultaneously or not. Making sure that this is the case would require to fit our mean model and residual dispersion models as a DHGLM.

### 4.4 Conclusion

The application of likelihood methods for mixed models to build and study isoscapes has been under-developed so far. In this paper we have introduced this approach to unfamiliar readers, but we have also detailed lesser known aspects and unusual adjustments of these methods to the present problems. As we have shown, our approach can be used to analysed the very large public dataset of the isotope composition of precipitation water provided by the Global Network for Isotopes in Precipitation (IAEA/WMO, 2017), but it can also be used to analyse other sources of data (*e.g.* the isotope composition of oceanic water data). For easier access to the methods, we have provided both a small `R` package called `ModellingIsoscapesUsingMixedModels` (included as Supplementary material) illustrating how to code the approach using the more general package `spaMM`, and the package `IsoriX` (Courtiol et al., 2016), which also uses `spaMM` internally, but which is dedicated to performing isoscape analyses and designed to be very simple to use. The latter package also allows for the inference of origin location of environmental materials using mixed models. All these packages are free and open source, similarly to the environment required to run them. By providing our approach in such a way, we allow for an efficient and objective comparison of different methods. We thus encourage other authors to make their code available and to compare alternative methodology for building isoscapes to ours. We also invite anyone interested to contribute to the development `IsoriX` of using GitHub (https://github.com/courtiol/IsoriX_project.

## Glossary

- Fixed effects describe the effects of known predictor variables and random effects describe the effects of unknown predictor variables. In the latter case, these effects are supposed to follow some probability distribution, which parameters are estimated.
- The residual error in a model is the difference between the response value and the expected value, given the realized value of the random effects. The estimates of the residuals are the difference between the response value and the predicted value. By construction, estimated residuals sum to zero in a linear model.
- LM, LMM: A function *f* is linear with respect to a vector-valued argument, ***β***, if *f* (***β*** = ***β***_1_ + ***β***_2_) = *f* (***β*** = ***β***_1_) + *f* (***β*** = ***β***_2_). A linear model (LM) is so called because the expected value of the response is linear with respect to the vector of fixed-effect coefficients. By definition, the distribution of the residual error in a LM is Gaussian. A linear mixed-effects model (LMM) is a model where the response, conditional on the realized values of the random effects, follows a LM, and the random effects have a Gaussian distribution.
- GLM, GLMM: A GLM is a model where the distribution of the residual error is not necessarily Gaussian, but rather belongs to the so-called exponential family of distributions, including the Gamma family considered in this work. In a GLM the expected value of the response may be a non-linear function of the estimated parameters, but is defined as a transformation (*e.g.* exponential) of a linear function (the linear predictor) of these parameters. This transformation is standardly specified by its inverse, the link (*e.g.* the log link for the exponential transformation). A GLM is fully specified by its linear predictor, family, link, and prior weights (see GLM specification). A generalized linear mixed-effect models (GLMM) is a model where the response, conditional on the realized values of the random effects, follows a GLM, and the random effects have a Gaussian distribution.
- GLM specification: Each distribution in the Gaussian and Gamma families is identified by its mean *μ* and its dispersion parameter *Φ*. The model for the mean is determined by the linear predictor and the link. The linear predictor is specified by the `formula` argument. The family and link are jointly specified by the `family` argument of the form *family(link)*, in particular `family=Gamma(link=log)` and `family=gaussian(link=identity)`(which can be entered as `family=gaussian()`). The model for *Φ* is by default a single estimated value for all levels of the response, but it can be modified by the *prior weights:* a vector of *prior weights* (*w*_*j*_) allows one to specify different variances for different levels of the response variable: for a given *Φ* the variance of the *j*th response is *Φ*/*w*_*j*_. Prior weights are given by the `prior.weight` argument in `spaMM`, which corresponds to the `weight` argument for `glm()`.

## ACKNOWLEDGEMENTS

We thank: Keith Hobson, Stephanie Kramer-Schadt, Marie-Sophie Rohwaeder, David Soto, Christian Voigt and Leonard Wassenaar for numerous discussions related to isotopes and isoscapes, as well as for feedbacks on this manuscript. This research did not receive any specific grant from funding agencies in the public, commercial, or not-for-profit sectors.

## Footnotes

1 For work including observations in the southern hemisphere, Bowen and Wilkinson (2002) recommend to use the absolute latitude. It makes no difference here as all latitudes we use are positive.

2 For *n* independent samples of a Gaussian distribution with variance σ^2^ and unknown mean, let *S*^2^ be the usual variance estimator (*i.e.* debiased by *n* — 1 denominator). Then (*n* − 1)*S*^2^/σ^2^ follows a chi-squared distribution with *n* − 1 degrees of freedom. This is a special case of the Gamma distribution with mean *n* − 1 and variance 2(*n* − 1). Then *S*^2^ is Gamma-distributed with mean σ^2^ and variance 2σ^2^/(*n* − 1).

3 Even though the absolute latitude can be used as a covariate for the fixed effect part of the model, the signed latitude should always be used in the Matérn term otherwise the computation of distances would be wrong.

4 The method is sometimes referred as estimating prediction errors in “*N* − 1 *jackknife*” (Bowen and Revenaugh, 2003).

5 Here, we do not provide units as the variation in estimates represent variations in logarithm of expected response, and such variations are dimensionless.

6 http://topotools.cr.usgs.gov/gmted_viewer/.

